# JAZF1 is transcriptionally regulated by SOX11 and promotes cardiac fibrosis by PI3K-Akt pathway

**DOI:** 10.1101/2023.09.01.555901

**Authors:** Yujing Mo, Rui Wang, Yingcong Liang, Yingling Zhou, Ying Zhang, Ling Xue

**Affiliations:** Department of Cardiology, Guangdong Cardiovascular Institute, Guangdong Provincial People’s Hospital (Guangdong Academy of Medical Sciences), Southern Medical University, Guangzhou, China

## Abstract

**Background:** Cardiac fibrosis is a component of all chronic heart diseases. JAZF1 regulates metabolism through various mechanisms; however, its role in cardiac fibrosis remains unclear. We aimed to investigate the role of JAZF1 in cardiac fibrosis.

**Methods:** A rat cardiac fibrosis model was established by administering isoproterenol subcutaneously for 14 days (5 mg/kg/day); an equal volume of saline was administered to the control group. Cardiac fibroblasts (CFs) were treated with TGF-β1 for 48 h to mimic cardiac fibrosis in vitro.

**Results:** JAZF1 expression at the protein and mRNA levels was enhanced in CFs and cardiac fibrosis tissues. JAZF1 downregulation suppressed CFs’ proliferation and migration. Western blotting showed that the PI3K/Akt signaling pathway was significantly decreased after JAZF1 knockdown. Further experiments revealed that SOX11 is an important transcription factor whose overexpression and downregulation enhanced and suppressed JAZF1 levels, respectively. Luciferase analysis showed that SOX11 interacted with the JAZF1 promoter. Moreover, SOX11 promoted cardiac fibrosis by regulating JAZF1 expression.

**Conclusions:** JAZF1 was enhanced in cardiac fibrosis tissue and TGF-β-treated CFs. JAZF1 knockdown decreased CFs’ migration and proliferation, possibly remediated by SOX11 with activation of PI3k/Akt signaling pathways.

## Introduction

Cardiac fibrosis is a component of all chronic heart diseases contributing to diastolic dysfunction and enhanced wall stiffness. Irrespective of the pathogenesis of cardiac fibrosis, activated fibroblasts regulate the development of fibrosis by producing excess extracellular matrix^1,2^. No specific therapy is currently available to stop the advancement of cardiac fibrosis, although several studies have demonstrated distinct antifibrotic strategies in proof-of-concept research^3,4^. Antifibrotic outcomes in ongoing clinical trials have been disappointing, with promising data coming mostly from therapies using renin-angiotensin-aldosterone system inhibitors^5^. Serelaxin is a reconstituted form of human relaxin-2, and Tim Wilhelmi discovered that serelaxin raises its receptor RXFP1 expression, which regulates the suppression of cardiac fibrosis and endothelial to mesenchymal transition; however, it fails to achieve its main endpoint as a treatment for acute heart failure in the clinical trials^6^. The design of effective therapies is impeded by the unclear underlying basis of cardiac fibrosis; therefore, studies examining the molecular mechanisms that participate in the progression of the disease are crucial.

JAZF1, a multifunctional regulator, was first established as a co-repressor of the orphan nuclear receptor (TR4)^7^. JAZF1, as a transcriptional co-regulator, can interact with protein kinases that participate in the metabolism of cellular energy and an array of nuclear receptors. Additionally, studies have indicated that JAZF1 participates in glucose metabolism, insulin sensitivity, cell differentiation, lipid metabolism, and inflammation^8–11^. JAZF1 regulates metabolism via different mechanisms, from suppressing inflammatory responses to enhancing glucose/lipid metabolism. However, its role in cardiac fibrosis remains unclear.

The sex-determining region Y-associated high-mobility group box transcription factor 11 (SOX11) is a critical member of the SOX transcription factor family and an essential regulator of embryonic development^12^. SOX11, along with SOX12 and SOX4, makes up the strongly conserved SOXC group^13^. Recent studies have revealed that SOX11 mRNA is among the most commonly elevated transcripts in many human cancers, including mantle cell lymphoma^14^, epithelial ovarian cancer^15^, breast cancer^16^, gastrointestinal tumors^17^, and nervous system neoplasms.

Moreover, SOX11 is considered a transcriptional activator in nervous system development. In patients with spinal cord injury, SOX11 can regulate the inflammatory response^18^. Furthermore, defective expression of SOX11 caused malformations in various human organs and systems, including the lungs, heart, and skeletal and gastrointestinal systems^19^. A recent study showed that SOX11 downregulation in fetal heart tissue mediated cardiomyocyte apoptosis^20^.

In the present study, we aimed to investigate the role of JAZF1 in cardiac fibrosis..

## Materials and Methods

### Ethics statement

All animal experiments were approved by the Ethics Committee of Guangdong Provincial Hospital(No.KY-Z-2021-309-02). Efforts were made to reduce pain in the animals.

### Animal grouping and model establishment

Twelve healthy specific pathogen-free male Sprague Dawley (SD) rats (200 ± 20 g) aged 6–8 weeks were acquired from Hunan SJA Laboratory Animal Co., Ltd [Hunan, China; license number: SCXK (xiang) 2019-0004]. After 1 week of acclimatization feeding, all animals were randomly allocated to the model and control groups (n = 6 each).

To establish the rat cardiac fibrosis model, isoproterenol (ISO) was administered subcutaneously for 14 days (5 mg/kg/day); an equal volume of saline was administered to the control group^21^. At the end of the 14 days, the rats were anesthetized using isoflurane (2.5% for induction and 1.0% for maintenance), and echocardiography was performed using the VisualSonics Vevo 2100 system(Visual Sonics, Amsterdam, Netherlands). Finally, to collect the organ, the rats were sacrificed by administering an overdose of sodium pentobarbital (75 mg/kg) intravenously; the hearts collected were used for molecular, pathological, and biological studies.

### Hematoxylin and eosin staining

Cardiac tissue was fixed in 4% paraformaldehyde, embedded in paraffin, sliced to a 4 μm thickness, dewaxed with dimethyl benzene, and dehydrated using gradient ethanol. Lastly, hematoxylin and eosin (H&E) staining was performed, and the sections were observed under a microscope.

### Masson staining

Heart tissue sections were degreased using xylene, rehydrated in graded alcohol, and stained using Masson’s Trichrome Staining kit (G1343, Solarbio, Beijing, China) following the manufacturer’s instructions. Next, the tissue sections were visualized under a microscope. The myocardium and fibrotic tissue were stained red and blue, respectively.

### Cell Counting Kit-8 assay

Cell viability was evaluated using Cell Counting Kit-8 (CCK-8) assay (Sangong Biotech, Shanghai, China) following the manufacturer’s protocol. Briefly, the cells were inoculated into 96-well microplates at a density of 2 × 10^3^ cells/well and cultivated in a 37°C humidified incubator at 5% CO_2_ for 1h. Finally, a spectrophotometer (BioTek, Winooski, VT, USA) was used to determine each well’s optical density at 450 nm.

### Migration assay

Migration assays were performed using 24-well Transwell plates (Corning Inc, Corning, NY, USA) per the manufacturer’s instructions. The cells were inoculated into the upper chamber of the plates at a density of 2 × 10^4^ cells/well without serum medium. The lower chamber was charged with 20% fetal bovine serum (FBS) medium (600 μL). After 24 h, the cells on the other side of the chamber were fixed with 4% formaldehyde and stained with crystal violet (Sangong Biotech).

### Quantitative real-time polymerase chain reaction

Total RNA was extracted from the cells using the TRIzol reagent (Invitrogen, Waltham, MA, USA) following the manufacturer’s instructions. Next, cDNA was synthesized using All-in-One First-Strand cDNA Synthesis SuperMix (TransGen Biotech, Beijing, China) for quantitative real-time polymerase chain reaction (qRT-PCR). To assess the mRNA levels of SOX11, JAZF1, IRF3, KLF4, and STAT5B, qRT-PCR was conducted using AceQ Universal SYBR qPCR Master Mix (Vazyme, Nanjing, China) on an RT-PCR system (Illumina Eco, California, USA). The primers used are listed in Table 1. Relative mRNA levels were calculated using the method of 2^−ΔΔCt^, taking GAPDH as an internal reference.

**Table 1.**
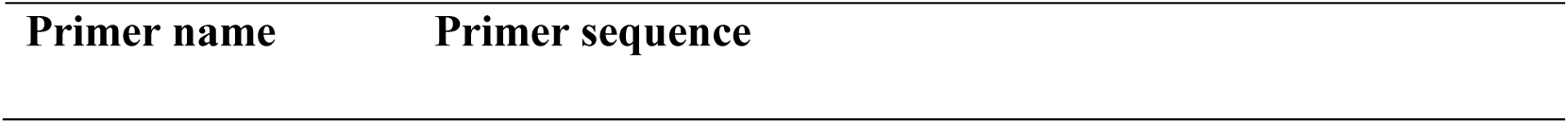
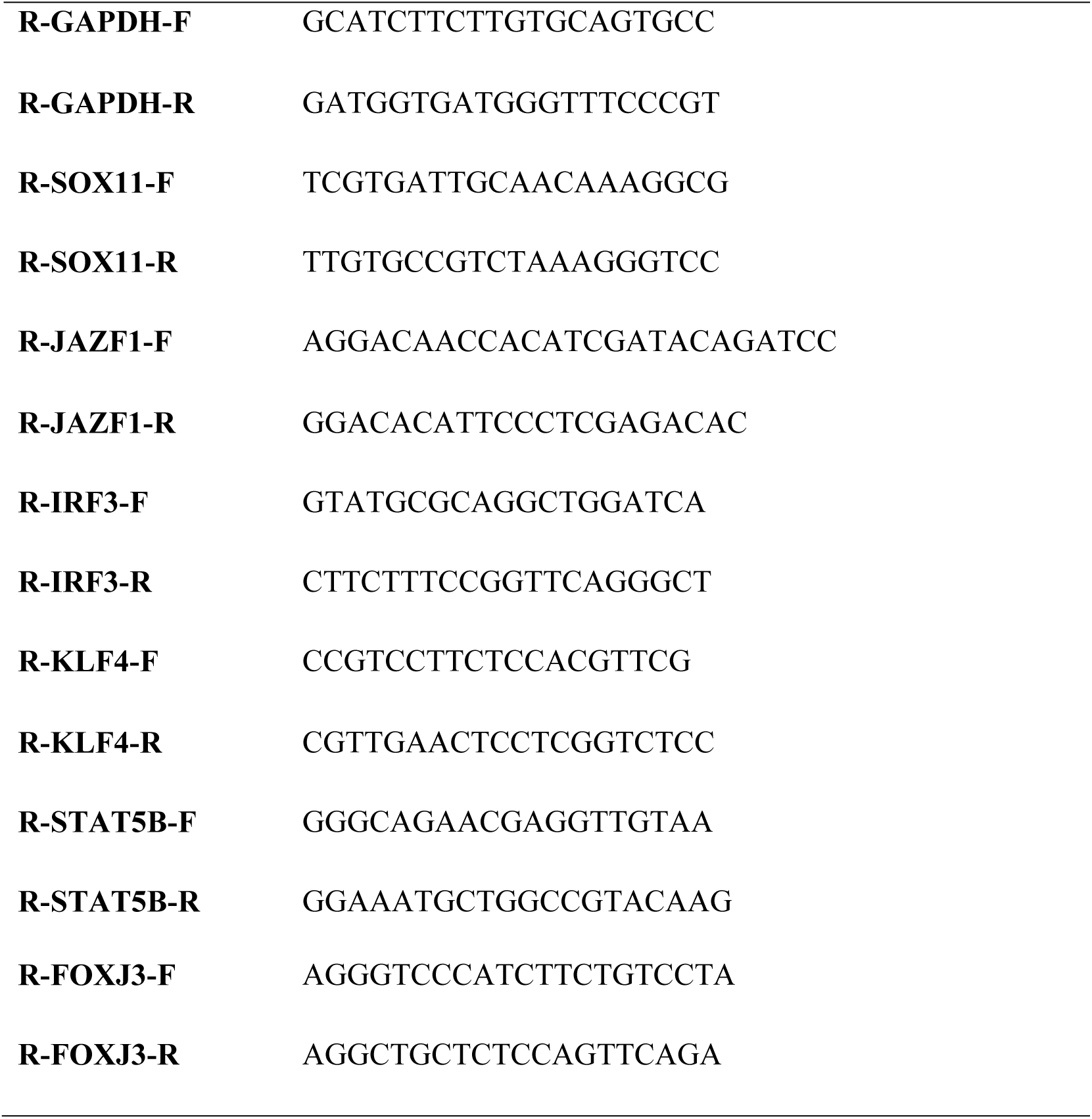
Sequences of the primers used for quantitative real-time polymerase chain reaction assays.

### Cell culture and treatments

Cardiac fibroblasts (CFs) were isolated from the hearts of neonatal SD rats (aged 0–3 days), inoculated in Dulbecco’s Modified Eagle Medium supplemented with 10% FBS, and cultivated in a 37°C cell culture incubator at 5% CO_2_. To mimic cardiac fibrosis in vitro, second-passaged CFs were injected into six-well plates, incubated until 70–80% confluence, and treated with the indicated concentration of TGF-β1(0ng/ml, 1ng/ml, 5ng/ml, 10ng/ml, 15ng/ml, 25ng/ml) for 48 h. For gene knockdown or overexpression, CFs were inoculated with siRNAs, plasmids, or lentiviruses in six-well plates and grown to 50–80% confluence. The transfection was performed using Lipofectamine 2000 (Invitrogen). After transfection for 24 h or infection with lentivirus for 48 h, the cells were treated with TGF-β1 for 48 h as needed.

### siRNAs, plasmid, and lentivirus

Rat JAZF1 and SOX11 siRNA were designed and synthesized by RiboBio (Guangzhou, China). The RNA sequences were as follows: GGACAACCACATCGATACA (JAZF1-siRNA1), CAAGAATGTGAACGGCATA (JAZF1-siRNA2), CACAATCAATTTCCATCCC (JAZF1-siRNA3), GGCTCTACTACAGCTTCAA (SOX11-siRNA1), GACCTGGTGTTCACGTATT (SOX11-siRNA2), and CCTGGACGACGATGATGAA (SOX11-siRNA3).

Two of the three siRNAs that effectively knocked down JAZF1 were used to construct lentivirus. The sequences were inserted into the NdeI/EcoRI restriction sites of the pLKO.1-Puro vector. Additionally, constructed plasmids and the packaging plasmids were transfected into HEK293 cells. After 48 hours, the supernatant containing the target lentivirus was collected. For SOX11 overexpression, the coding sequence of rat SOX11 was obtained using PCR and subcloned into the pCDH-GFP+Puro vector to construct overexpression plasmids.

### Western blotting

Proteins in the rat tissues or cells were extracted using RIPA lysis buffer (Beyotime Biotechnology, Shanghai, China). Next, equal quantities of proteins were separated using sodium dodecyl sulfate-polyacrylamide gel electrophoresis and transferred to the polyvinylidene difluoride membranes, which were incubated at 4°C overnight with the primary antibodies that were diluted in bovine serum albumin. The primary antibodies were α-SMA (A5228, Sigma-Aldrich, St. Louis, MI, USA), collagen I (ab260043, Abcam, Cambridge, UK), collagen III (22734-1-AP, Proteintech, Rosemont, IL, USA), SOX11 (ab234996, Abcam), JAZF1(sc-376503, Santacruz, Dallas, TX, USA), GAPDH(#5174, CST, Denver, MA, USA), PI3K(ab191606, Abcam, p-AKT(4060, CST), AKT(4691, CST), and P-PI3K (ab182651, Abcam). Moreover, secondary antibodies,anti-mouse, and anti-rabbit IgG were labeled using horseradish peroxidase (CST) and diluted in 5% TBST; the PVDF membranes were inoculated with secondary antibodies at 37°C for 120 min. Finally, an electrochemiluminescence substrate (Thermo Fisher Scientific, Waltham, MA, USA) was used to visualize the immunoreactive bands exposed to an X-ray film (SUPER RX-N-C, China).

### Dual-luciferase reporter assay

Luciferase activity was investigated using the Dual-Luciferase® Reporter Assay System (E1910, Promega, Madison, WI, USA) per the manufacturer’s instructions. Briefly, firefly luciferase reporter plasmids carrying the JAZF1 promoter were constructed and co-transfected with SOX11 overexpressing or vector plasmids into CFs. Finally, the relative luciferase activity was calculated as follows:

Relative luciferase activity = firefly luciferase activity/Renilla luciferase activity.

### Statistical analysis

All data are presented as the means ± standard deviation for triplicate determinations. The statistical significance between the model and control groups was estimated using the student’s t-test. Additionally, multiple comparisons were performed using one- or two-way analysis of variance. GraphPad Prism version 9.0 (GraphPad Software, San Diego, CA, USA) was used for all the analyses. Statistical significance was set at P < 0.05.

## Results

### JAZF1 expression was enhanced during cardiac fibrosis

To determine the underlying role of JAZF1 in the progression of cardiac fibrosis, we established an ISO-induced rat model of cardiac fibrosis. Cardiac fibrosis was evaluated using Masson’s trichrome indicators, H&E staining, and echocardiography. Echocardiography revealed that the ISO-induced group had decreased ventricular systolic function (ejection fraction), fractional shortening, and ventricular dilatation compared with the control group (Figs. 1A–B). Masson’s trichrome and H&E staining revealed that the fibrosis area increased sharply with exposure to ISO (Fig. 1C). Additionally, protein levels of fibrosis biomarkers, α-SMA, collagen I, and collagen III, were elevated in the ISO-induced cardiac fibrosis tissue compared with the control group (Fig. 1D). These results indicate the successful construction of a rat cardiac fibrosis model. JAZF1 expression was subsequently detected; JAZF1 protein levels were upregulated in cardiac fibrosis.

**Figure 1:**
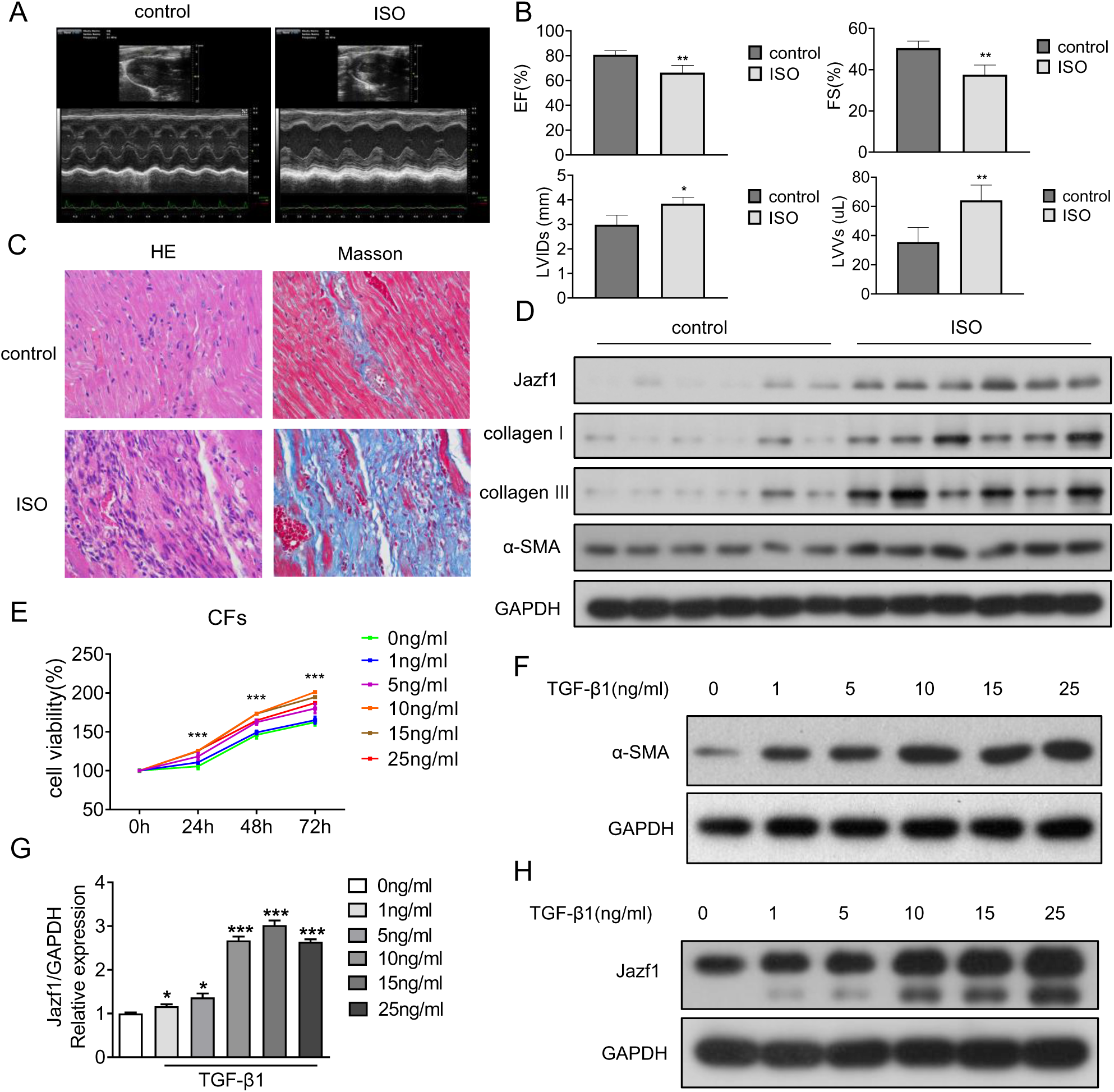
JAZF1 expression was enhanced during cardiac fibrosis. A-B: Ventricular systolic function (ejection fraction), fractional shortening, and ventricular diameter by echocardiography of each group. C: Masson’s trichrome and H&E staining of each group. D: Western blotting of α-SMA, collagen I, and collagen III in control group and ISO group. E-F: CCK-8 assay and western blotting revealed cell viability and α-SMA expression, respectively, with the increasing concentration of TGF-β1. G-H: JAZF1 protein and mRNA levels in TGF-β1-treated CFs. α-SMA: Smooth muscle alpha-actin, TGF-β1: transforming growth factor beta 1; ISO: isoproterenol; * *P*<0.05; ** *P*<0.01; *** *P*<0.001, Twelve healthy specific pathogen-free male Sprague Dawley (SD) rats were included and randomly allocated to the model and control groups (n = 6 each).

To examine the role of JAZF1 in CFs in vitro, we isolated cardiac fibroblasts from SD rats and treated them with different concentrations of TGF-β1 to mimic a cellular cardiac fibrosis model. CCK-8 assay and western blotting revealed increased cell viability and α-SMA expression, respectively, with the increasing concentration of TGF-β1 (Figs. 1E–F). Lastly, JAZF1 protein and mRNA levels were raised in TGF-β1-treated CFs (Figs. 1G–H).

### JAZF1 knockdown prevented TGF-β1-induced cardiac fibrosis

To gain insight into the role of JAZF1 in cardiac fibrosis, a negative control siRNA and three siRNAs targeting JAZF1 were designed and transfected into CFs. The knockdown efficiency was confirmed using qRT-PCR and western blotting (Figs. 2A and B). JAZF1 expression silencing significantly suppressed CFs’ viability, migration, and fibrosis with or without TGF-β1 treatment (Figs. 2C and D). The Levels of the representative cardiac fibrosis markers including collagenI, collagenIII and α-SMA in CFs induced by TGF-β1 were reversed by downregulation of JAZF1 (Fig. 2E).

**Figure 2:**
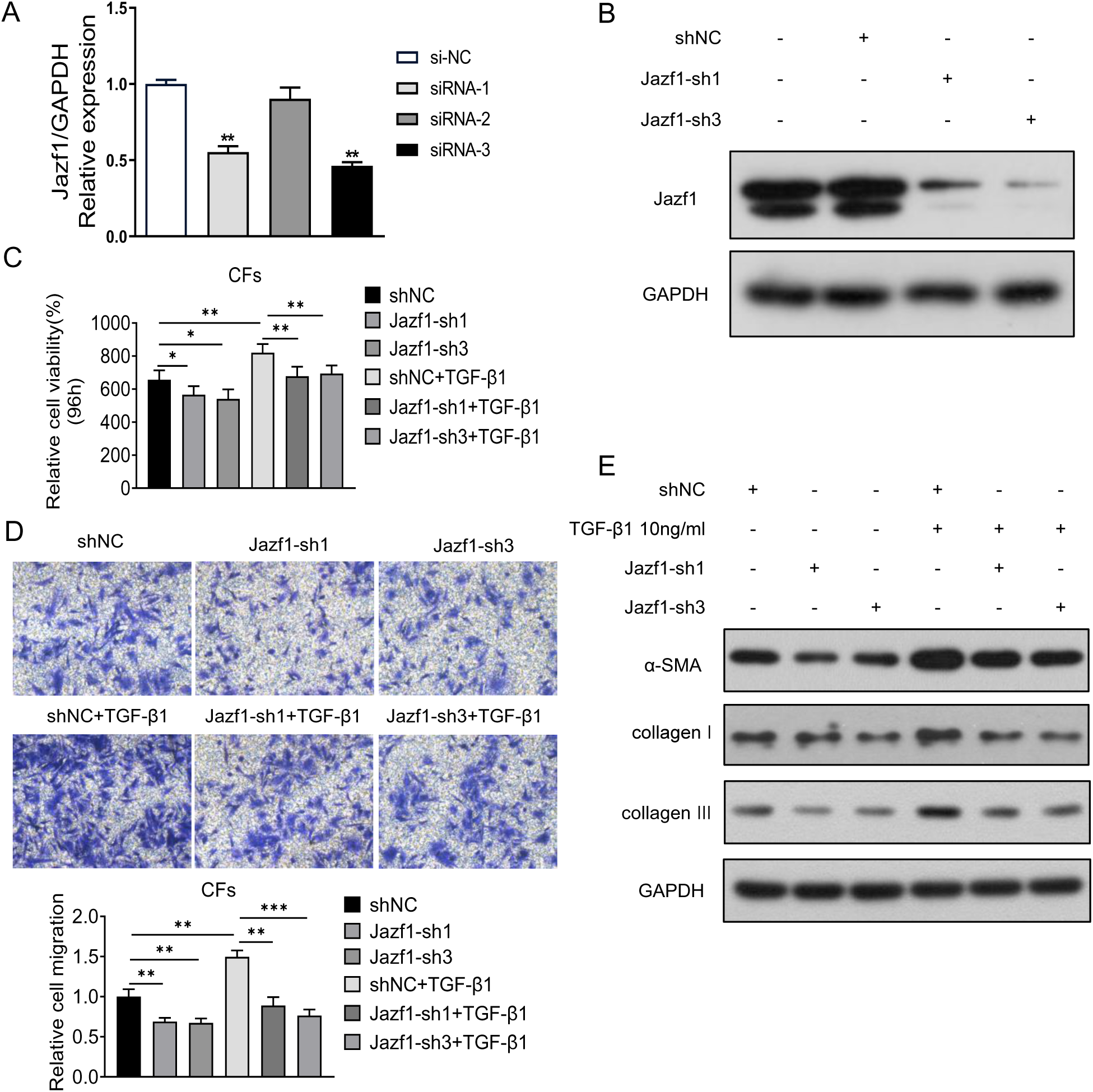
JAZF1 knockdown prevented TGF-β1-induced cardiac fibrosis. A-B: The RNA and protein level of JAZF1 in JAZF1 knockdown cells. C-D: Cell viability, fibrosis and cell migration of JAZF1 knockdown CFs with and without TGF-β1 treatment. E: Collagen I, collagen III and α-SMA in CFs induced by TGF-β1 with and without downregulation of JAZF1 TGF-β1: transforming growth factor beta 1; GAPDH: glyceraldehyde-3-phosphate dehydrogenase; ISO: isoproterenol; CFs: Cardiac fibroblasts; sh: small hairpin RNAs; NC: none; * *P*<0.05; ** *P*<0.01; *** *P*<0.001.

### JAZF1 regulated PI3K/Akt pathway in CFs

To clarify the possible mechanisms of JAZF1 in cardiac fibrosis, we performed RNA sequencing of CFs with and without JAZF1 knockdown. We obtained 506 differentially expressed genes by comparing the expression profiles (Fig. 3A), which were analyzed using the KEGG pathway. KEGG pathway enrichment was visualized using a bubble diagram, in which we noted the PI3K/Akt pathway (Fig. 3B). The PI3K/Akt pathway has been extensively associated with fibrosis, including cardiac fibrosis ^22–24^. Therefore, a western blot assay was conducted to explore whether the PI3K/Akt pathway was involved in JAZF1’s role in CFs. PI3K and Akt phosphorylation decreased in JAZF1 siRNA-transfected cells (Fig. 3).

**Figure 3:**
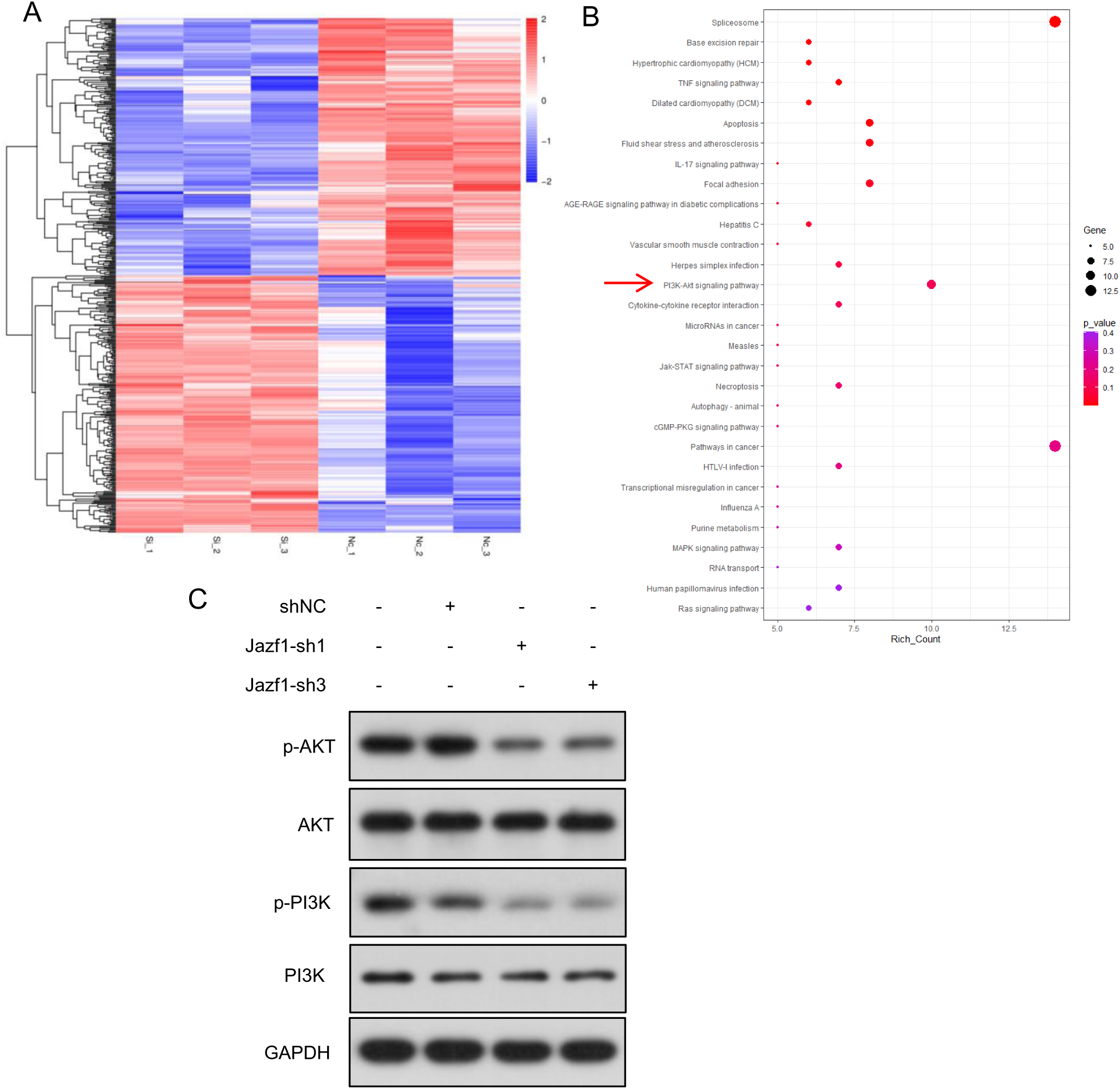
Jazf1 regulates myocardial fibrosis through PI3K-Akt pathway. A: Heat map of differentially expressed genes after JAZF1 knockdown in CF cells; B) KEGG enrichment pathway; C) PI3K-Akt pathway changes after Jazf1 knockdown. CFs: Cardiac fibroblasts; GAPDH: glyceraldehyde-3-phosphate dehydrogenase; sh: Small hairpin RNAs; SiRNA: Small interfering RNA; NC: none; p-AKT: phosphorylated AKT; p-PI3K: phosphorylated PI3K; * *P*<0.05; ** *P*<0.01; *** *P*<0.001.

### JAZF1 was transcriptionally activated by SOX11

To explore the drivers of upregulated JAZF1 expression during cardiac fibrosis, potential transcription factors of JAZF1 were predicted using JASPAR (https://jaspar.genereg.net/). Five candidates with high scores were selected for the experimental validation. Two (SOX11 and IRF3) out of five candidate transcription factors were significantly upregulated in TGF-β1 induced CFs (Figs. 4A–E); however, SOX11 experienced the largest change. Thus, we studied the SOX11 regulatory effect on JAZF1 and its effects on cardiac fibrosis. Using tissues from a previously prepared rat cardiac fibrosis model, SOX11 upregulation during cardiac fibrosis was confirmed in vivo using western blotting (Fig. 4F). Moreover, SOX11 overexpression significantly enhanced the mRNA and protein levels of JAZF1 (Figs. 4G and H); however, SOX11 knockdown suppressed JAZF1 expression (Fig. 4I).

**Figure 4:**
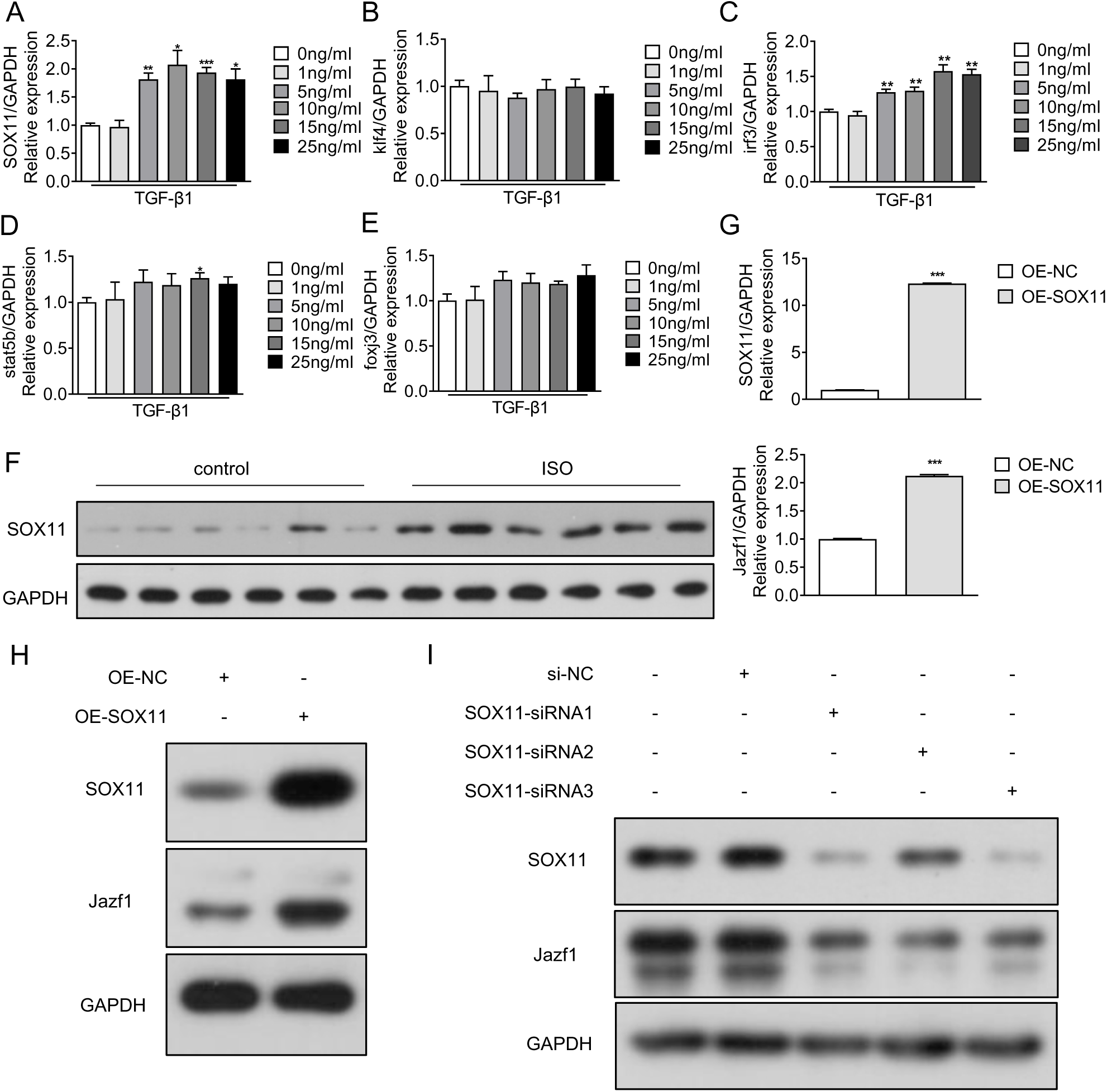
JAZF1 was transcriptionally activated by SOX11. A-E: Five candidate transcription factors’ expression in TGF-β1 induced CFs; F: SOX11 expression in CFs; G: RNA level of SOX11 change with JAZF1; H: JAZF1 protein change with SOX11 overexpression; I: Protein level of JAZF1 after knocking down SOX11; GAPDH: glyceraldehyde-3-phosphate dehydrogenase; ISO: isoproterenol; OE: overexpression; NC: None; SiRNA: Small interfering RNA; CFs: Cardiac fibroblasts; * *P*<0.05; ** *P*<0.01; *** *P*<0.001.

To confirm the direct interaction between SOX11 and JAZF1 promoters, we performed a dual-luciferase assay; SOX11 increased the luciferase activity of JAZF1 reporter plasmids (Fig. 5A). To gain further insight into the regulation of cardiac fibrosis by JAZF1, we investigated whether JAZF1 is responsible for CFs’ behavior in the presence of SOX11. CFs’ viability, migration, and fibrosis were notably increased by SOX11 overexpression and reversed by JAZF1 knockdown (Figs. 5B–D).

**Figure 5:**
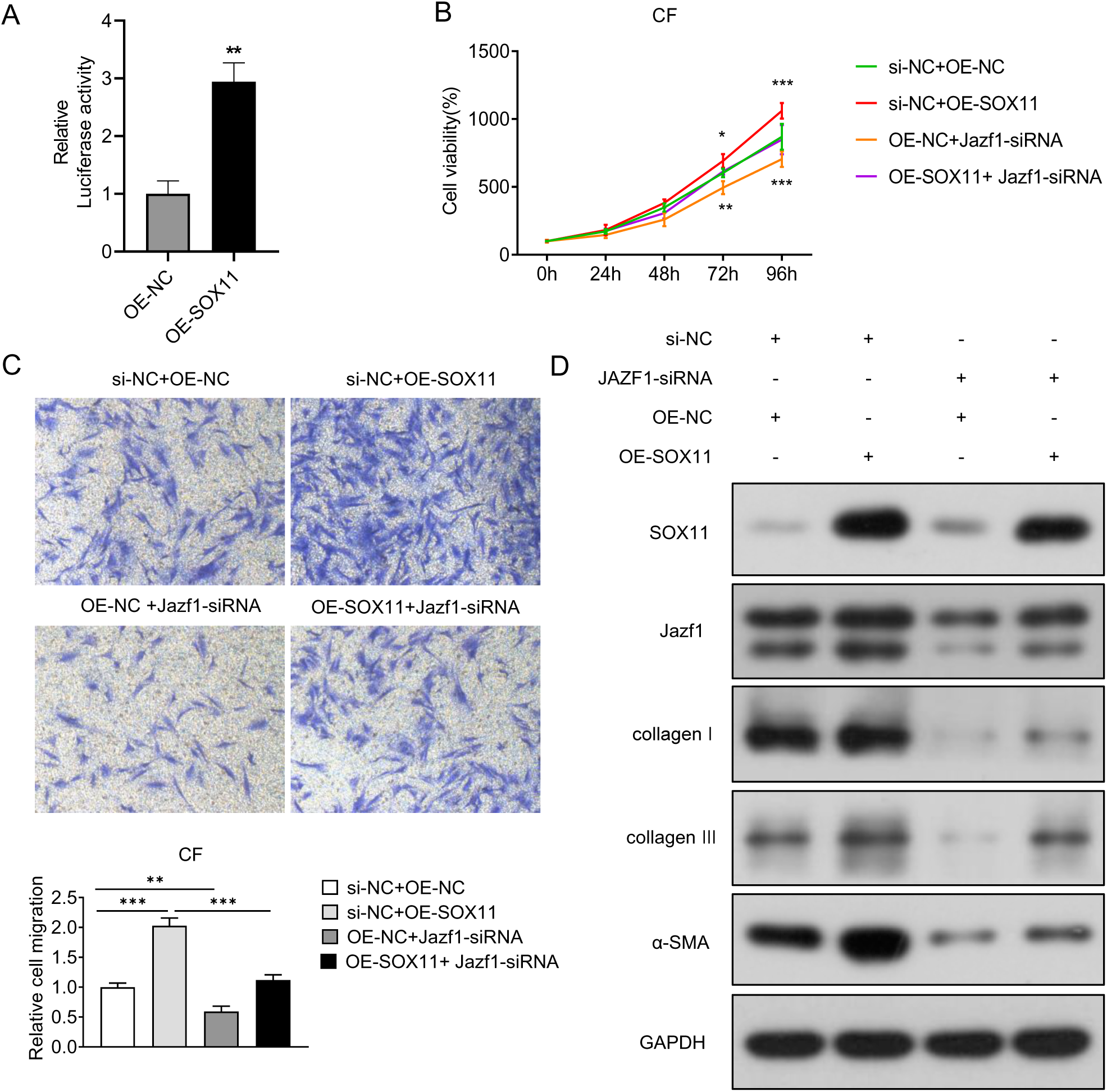
JAZF1 was transcriptionally activated by SOX11. A: Luciferase assay verified the promoter binding between SOX11 and JAZF1; B-D: JAZF1’s response to SOX11, in CFs’ viability, migration, and fibrosis markers. OE: overexpression; NC: None; SiRNA: Small interfering RNA; CFs: Cardiac fibroblasts; GAPDH: glyceraldehyde-3-phosphate dehydrogenase; * *P*<0.05; ** *P*<0.01; *** *P*<0.001.

## Discussion

Myocardial fibrosis is characterized by various qualitative and quantitative alterations in the interstitial collagen network of the myocardium following ischemic injury to the heart from drugs, systemic disease, or any other deleterious stimulus affecting the heart or circulatory system. Therefore, cardiac fibrosis is the primary driver of the increasing burden of heart failure, particularly in the elderly^25^. However, cardiac fibrosis inclusion in heart failure management remains an unmet medical need. Developing more specific in vivo methodologies and recognizing new biomarkers will permit better investigation of cardiac fibrosis. This study suggests that JAZF1 plays a key role in cardiac fibrosis; its overexpression causes the migration and proliferation of CFs, which are important mediators of cardiac fibrosis ^1^. Moreover, in this study, d JAZF1 was overexpressed in cardiac fibrosis tissue and TGF-β-treated CFs, and its downregulation reversed fibrosis in CFs. Therefore, JAZF1 overexpression caused cardiac fibrosis.

JAZF1 encodes a nuclear protein containing three C2H2-type zinc fingers and is a transcriptional repressor. Additionally, JAZF1 is hypothesized to act as a tumor suppressor, particularly in prostate cancer and endometrial stromal sarcomas^26,27^. Previous studies suggested that JAZF1 is involved in lipid metabolism, insulin sensitivity, inflammation, and glucose metabolism^9^. Furthermore, Yuan et al.^10^ discovered that JAZF1 plays a major role in modulating insulin sensitivity and liver glucose metabolism during glucose homeostasis by activating the Akt signaling pathway. Kobiita et al.^28^ showed that JAZF1 is a transcriptional regulator transferred to the nucleus during diabetes and metabolic stress. Additionally, they demonstrated some functions of JAZF1 in modulating genes that participate in ribosome biogenesis and protein translation, including the transcription of insulin genes. The role of JAZF1in heart disease has seldom been reported before. Bae et al. ^29^ found that JAZF1-overexpressing transgenic mice experienced myocardial apoptosis, leading to mitochondrial defects, ECG abnormalities, and high blood pressure. As far as we know, this is the first study exploring the effect of JAZF1 in cardiac fibrosis.

The PI3K/Akt pathway is one of the most critical intracellular signaling pathways. PI3K and Akt kinases were identified by Whitman M et al.^30^ in 1988 and Staal ^31^ in 1987, respectively. PI3K family members are proto-oncogenes, important kinases of inositol and phosphatidyl-inositol (PI), and important signal transduction molecules in cells involved in the modulation of apoptosis, proliferation and differentiation of cells, and other processes, such as phosphorylation of the 3’ hydroxyl on the PI ring^32^. Yang et al. ^33^ found that when pretreated with CpG-ODN, ISO-treated mice exhibited an isoproterenol-resistant phenotype, including a reduction in cardiac cell fibrosis and death. Moreover, in the heart, the expression levels of PI3Kγ were elevated in pathological states and were implicated in the remodeling of the heart, a key process in cardiac fibrosis^34^.

Akt is the central hub of signal transduction that modulates cellular functions from growth to survival and metabolism^35^. Research suggests that Members of the Akt family are expressed in the myocardium and play overlapping but diverse roles^22^. Furthermore, cumulative studies have demonstrated that Akt stabilizes the growth and proliferation of cells and suppresses apoptosis through some proteins and that the proliferation and differentiation of cardiac fibroblasts are essential processes in the progression of cardiac fibrosis. Aikawa et al.^36^ found that PI3K/Akt can phosphorylate and inactivate Bad, a key pro-apoptotic factor, thereby facilitating cell survival and blocking apoptosis. Lastly, Akt modulated the apoptotic pathway in cardiomyocytes through one of its substrates, caspase-9, a member of the caspase family of proteases^37^.

As stated, many studies have proven that PI3K/Akt signaling pathway is involved in many processes of cardiac fibrosis^22^. In our study, we confirmed JAZF1 participation in the PI3K/Akt signaling pathway using western blotting. After the JAZF1 knockdown, the PI3K/Akt signaling pathway was significantly decreased. These findings suggest that activation of the PI3K/Akt signaling pathway is implicated in the effect of JAZF1 on the migration and proliferation of CFs.

Furthermore, we examined how JAZF1 regulation affects cardiac fibrosis in CFs. The SOX gene family comprises numerous supergenes with a conserved high-mobility group frame pattern and is a critical transcriptional regulator^38^. SOX11 belongs to subgroup C of the SOX protein family. Aberrant modulation of SOX11 has been observed in many solid tumors, including epithelial ovarian tumors, gliomas, and medulloblastomas^39^. Moreover, SOX11 is reportedly essential for the pulmonary inflammatory response^40^, and SOX11 knockout mice demonstrated serious developmental defects, including defects related to the heart, spleen generation, skeletal development, and the development of the anterior ophthalmic ganglion^41^. Lastly, Su et al.^20^ identified that SOX11 overexpression represses apoptosis in cardiomyocytes after hyperglycemia treatment. In our study, SOX11 was upregulated in cardiac fibrosis samples. Additionally, SOX11 overexpression promoted CFs’ cell migration, proliferation, and fibrosis by transcriptionally regulating JAZF1, and JAZF1 knockdown blocked these promoting effects.

Summarily, JAZF1 expression was enhanced in cardiac fibrosis tissue and TGF-β1-treated CFs. Additionally, JAZF1 knockdown suppressed the CFs’ migration, proliferation, and fibrosis, which might be induced by the SOX11 upstream and PI3k/Akt signaling pathways downstream. Overall, our results indicate that JAZF1 may play a significant role in cardiac fibrosis. This study help us to understand the role of SOX11-JAZF1-PI3k/Akt in cardiac fibrosis and may contribute to the design of further studies related to this pathway, as well as sought to shed light on a potential treatment for cardiac fibrosis.

## Acknowledgments

None.

## Funding

This research was supported by the Natural Science Foundation of Guangdong Province (grant No. 2023A1515010177) and National Natural Science Foundation of China (grant No. 82000057).

## Declaration

### Competing interests

The authors declare no competing interests.

### Ethics approval and consent to participat

The experimental protocol was established, according to the ethical guidelines of laboratory animal and was approved by the Laboratory Animal Technology Ethics Committee of Guangdong Provincial People’s Hospital.

### Consent for publication

Not applicable.

### Availability of data and materials

All data generated or analyzed during this study are included in this published article.

### Authors’ contributions

Study conception and design: Yujing Mo, Rui Wang, Yingcong Liang and Ling Xue; experiment conduction: Yujing Mo, Rui Wang, Yingcong Liang and Ying Zhang; analysis and interpretation of results: Yujing Mo, Rui Wang, Yingling Zhou and Ling Xue; draft manuscript preparation: Yujing Mo, Yingling Zhou and Ling Xue. All authors reviewed the results and approved the final version of the manuscript.

## Reference

1. Krenning G, Zeisberg EM, Kalluri R. The origin of fibroblasts and mechanism of cardiac fibrosis. J Cell Physiol 2010;225:631–7.

2. Tallquist MD, Molkentin JD. Redefining the identity of cardiac fibroblasts. Nat Rev Cardiol 2017;14:484–91.

3. Tao L, Bei Y, Chen P, et al. Crucial role of miR-433 in regulating cardiac fibrosis. Theranostics 2016;6:2068–83.

4. Chen Z, Li Y, Dian K, Rao L. Modulating microRNAs as novel therapeutic targets in cardiac fibrosis. Theranostics 2017;7:2287–8.

5. Fang L, Murphy AJ, Dart AM. A clinical perspective of anti-fibrotic therapies for cardiovascular disease. Front Pharmacol 2017;8:186.

6. Wilhelmi T, Xu X, Tan X, et al. Serelaxin alleviates cardiac fibrosis through inhibiting endothelial-to-mesenchymal transition via RXFP1. Theranostics 2020;10:3905–24.

7. Adams MD, Soares MB, Kerlavage AR, Fields C, Venter JC. Rapid cDNA sequencing (expressed sequence tags) from a directionally cloned human infant brain cDNA library. Nat Genet 1993;4:373–80.

8. Johnson JA, Watson JK, Nikolić MZ, Rawlins EL. Fank1 and JAZF1 promote multiciliated cell differentiation in the mouse airway epithelium. Biol Open 2018;7:bio033944.

9. Meng F, Lin Y, Yang M, et al. JAZF1 inhibits adipose tissue macrophages and adipose tissue inflammation in diet-induced diabetic mice. Biomed Res Int 2018;2018:4507659.

10. Yuan L, Luo X, Zeng M, et al. Transcription factor TIP27 regulates glucose homeostasis and insulin sensitivity in a PI3-kinase/Akt-dependent manner in mice. Int J Obes (Lond) 2015;39:949–58.

11. Ming GF, Xiao D, Gong WJ, et al. JAZF1 can regulate the expression of lipid metabolic genes and inhibit lipid accumulation in adipocytes. Biochem Biophys Res Commun 2014;445:673–80.

12. Yang Z, Jiang S, Lu C, et al. SOX11: friend or foe in tumor prevention and carcinogenesis. Ther Adv Med Oncol 2019;11:1758835919853449.

13. Jay P, Gozé C, Marsollier C, et al. The human SOX11 gene: cloning, chromosomal assignment and tissue expression. Genomics 1995;29:541–5.

14. Wasik AM, Lord M, Wang X, et al. SOXC transcription factors in mantle cell lymphoma: the role of promoter methylation in SOX11 expression. Sci Rep 2013;3:1400.

15. Sernbo S, Gustavsson E, Brennan DJ, et al. The tumour suppressor SOX11 is associated with improved survival among high grade epithelial ovarian cancers and is regulated by reversible promoter methylation. BMC Cancer 2011;11:405.

16. Shepherd JH, Uray IP, Mazumdar A, et al. The SOX11 transcription factor is a critical regulator of basal-like breast cancer growth, invasion, and basal-like gene expression. Oncotarget 2016;7:13106–21.

17. Xu X, Chang X, Li Z, et al. Aberrant SOX11 promoter methylation is associated with poor prognosis in gastric cancer. Cell Oncol (Dordr) 2015;38:183–94.

18. Feng JS, Sun JD, Wang XD, Fu CH, Gan LL, Ma R. MicroRNA-204-5p targets SOX11 to regulate the inflammatory response in spinal cord injury. Eur Rev Med Pharmacol Sci 2019;23:4089–96.

19. Wurm A, Sock E, Fuchshofer R, Wegner M, Tamm ER. Anterior segment dysgenesis in the eyes of mice deficient for the high-mobility-group transcription factor Sox11. Exp Eye Res 2008;86:895–907.

20. Su D, Gao Q, Guan L, et al. Downregulation of SOX11 in fetal heart tissue, under hyperglycemic environment, mediates cardiomyocytes apoptosis. J Biochem Mol Toxicol 2021;35:e22629.

21. Tao H, Shi P, Zhao XD, Xuan HY, Ding XS. MeCP2 inactivation of LncRNA GAS5 triggers cardiac fibroblasts activation in cardiac fibrosis. Cell Signal 2020;74:109705.

22. Qin W, Cao L, Massey IY. Role of PI3K/Akt signaling pathway in cardiac fibrosis. Mol Cell Biochem 2021;476:4045–59.

23. Wang J, Hu K, Cai X, et al. Targeting PI3K/AKT signaling for treatment of idiopathic pulmonary fibrosis. Acta Pharm Sin B 2022;12:18–32.

24. Guo J. Effect of miR-21 on renal fibrosis induced by nano-SiO₂ in diabetic nephropathy rats via PTEN/AKT pathway. J Nanosci Nanotechnol 2021;21:1079–84.

25. Conrad N, Judge A, Tran J, et al. Temporal trends and patterns in heart failure incidence: a population-based study of 4 million individuals. Lancet 2018;391:572–80.

26. Prokunina-Olsson L, Fu YP, Tang W, et al. Refining the prostate cancer genetic association within the JAZF1 gene on chromosome 7p15.2. Cancer Epidemiol Biomarkers Prev 2010;19:1349–55.

27. Hrzenjak A. JAZF1/SUZ12 gene fusion in endometrial stromal sarcomas. Orphanet J Rare Dis 2016;11:15.

28. Kobiita A, Godbersen S, Araldi E, et al. The diabetes gene JAZF1 Is Essential for the HOMEOSTATIC Control of Ribosome Biogenesis and Function in Metabolic Stress. Cell Rep 2020;32:107846.

29. Bae KB, Kim MO, Yu DH, et al. Overexpression of JAZF1 induces cardiac malformation through the upregulation of pro-apoptotic genes in mice. Transgenic Res 2011;20:1019–31.

30. Whitman M, Downes CP, Keeler M, Keller T, Cantley L. Type I phosphatidylinositol kinase makes a novel inositol phospholipid, phosphatidylinositol-3-phosphate. Nature 1988;332:644–6.

31. Staal SP. Molecular cloning of the akt oncogene and its human homologues AKT1 and AKT2: amplification of AKT1 in a primary human gastric adenocarcinoma. Proc Natl Acad Sci U S A 1987;84:5034–7.

32. Naga Prasad SV, Perrino C, Rockman HA. Role of phosphoinositide 3-kinase in cardiac function and heart failure. Trends Cardiovasc Med 2003;13:206–12.

33. Yang L, Cai X, Liu J, et al. CpG-ODN attenuates pathological cardiac hypertrophy and heart failure by activation of PI3Kα-Akt signaling. PLoS One 2013;8:e62373.

34. Perino A, Ghigo A, Ferrero E, et al. Integrating cardiac PIP3 and cAMP signaling through a PKA anchoring function of p110γ. Mol Cell 2011;42:84–95.

35. Hers I, Vincent EE, Tavaré JM. Akt signalling in health and disease. Cell Signal 2011;23:1515–27.

36. Aikawa R, Nawano M, Gu Y, et al. Insulin prevents cardiomyocytes from oxidative stress-induced apoptosis through activation of PI3 kinase/Akt. Circulation 2000;102:2873–9.

37. Abeyrathna P, Su Y. The critical role of Akt in cardiovascular function. Vascul Pharmacol 2015;74:38–48.

38. Kavyanifar A, Turan S, Lie DC. SoxC transcription factors: multifunctional regulators of neurodevelopment. Cell Tissue Res 2018;371:91–103.

39. Vegliante MC, Royo C, Palomero J, et al. Epigenetic activation of SOX11 in lymphoid neoplasms by histone modifications. PLoS One 2011;6:e21382.

40. Mitamura Y, Nunomura S, Nanri Y, et al. Hierarchical control of interleukin 13 (IL-13) signals in lung fibroblasts by STAT6 and SOX11. J Biol Chem 2018;293:14646–58.

41. Paul MH, Harvey RP, Wegner M, Sock E. Cardiac outflow tract development relies on the complex function of Sox4 and Sox11 in multiple cell types. Cell Mol Life Sci 2014;71:2931–45.

